# A fast method to distinguish between fermentative and respiratory metabolisms in single yeast cells

**DOI:** 10.1101/2023.06.23.546324

**Authors:** L. Luzia, J. Battjes, V. E. Zwering, D. B. Jansen, C. Melkonian, B. Teusink

## Abstract

*Saccharomyces cerevisiae* adapts its metabolism according to nutrient availability. Typically, it rapidly ferments glucose to ethanol, and then shifts to respiration when glucose becomes limited. However, our understanding of the regulation of metabolism is largely based on population averages, whereas nutrient transitions may cause heterogeneous responses at the individual cell level. Although protein expression can be followed at the single-cell level as a proxy for metabolic modes, direct assessment of the contribution of respiration or (respiro)fermentation to energy metabolism is lacking. Here we describe a method to quickly differentiate between fermentative and respiratory metabolisms in individual cells of budding yeast. The method explores the use of the fluorescent FRET-based biosensor yAT1.03 to measure cytosolic ATP, coupled with the respiratory inhibitor Antimycin A. For the method validation, we used cells under fermentative and respiratory regimes from batch and chemostat cultures. Upon Antimycin A addition, we observed a sharp decrease of the normalized FRET ratio for respiratory cells; respirofermentative cells showed no response. Next, we tracked the changes in metabolism during the diauxic shift of a glucose pre-grown batch culture. Following glucose exhaustion, the entire cell population experienced a progressive rise in intracellular ATP produced via respiration, suggesting a uniform and gradual increase in respiratory capacity as cells pick up growth in a medium with ethanol as the sole carbon source. Overall, the combination of yAT1.03 with Antimycin A is a robust tool to distinguish fermentative from respiratory yeast cells, offering a new single-cell opportunity to study yeast metabolism.

**Graphical abstract:** Identification of fermentative and respiratory metabolisms in yeast cells using an ATP sensor coupled with a respiration inhibitor.
(a) yAT1.03 consists of a donor (tdTomato) and an acceptor (ymTq2Δ11) domain linked by a binding domain with affinity to ATP. When ATP binds to the binding domain, donor and acceptor come together and the Förster energy is transferred from the first to the second domain. When expressed in *in vivo* cells the sensor allows real time measurements of ATP changes. (b) Depending on the growth conditions, yeast cells expressing yAT1.03 show a distinct response after being pulsed with the respiratory inhibitor Antimycin A (AA). The drop in ATP levels in respiratory cells caused by AA results from the inhibition of the mitochondrial electron transport chain. (c) Distinct metabolic responses to an AA pulse pre-, during and post-diauxic shift reveal distinct metabolic phenotypes.

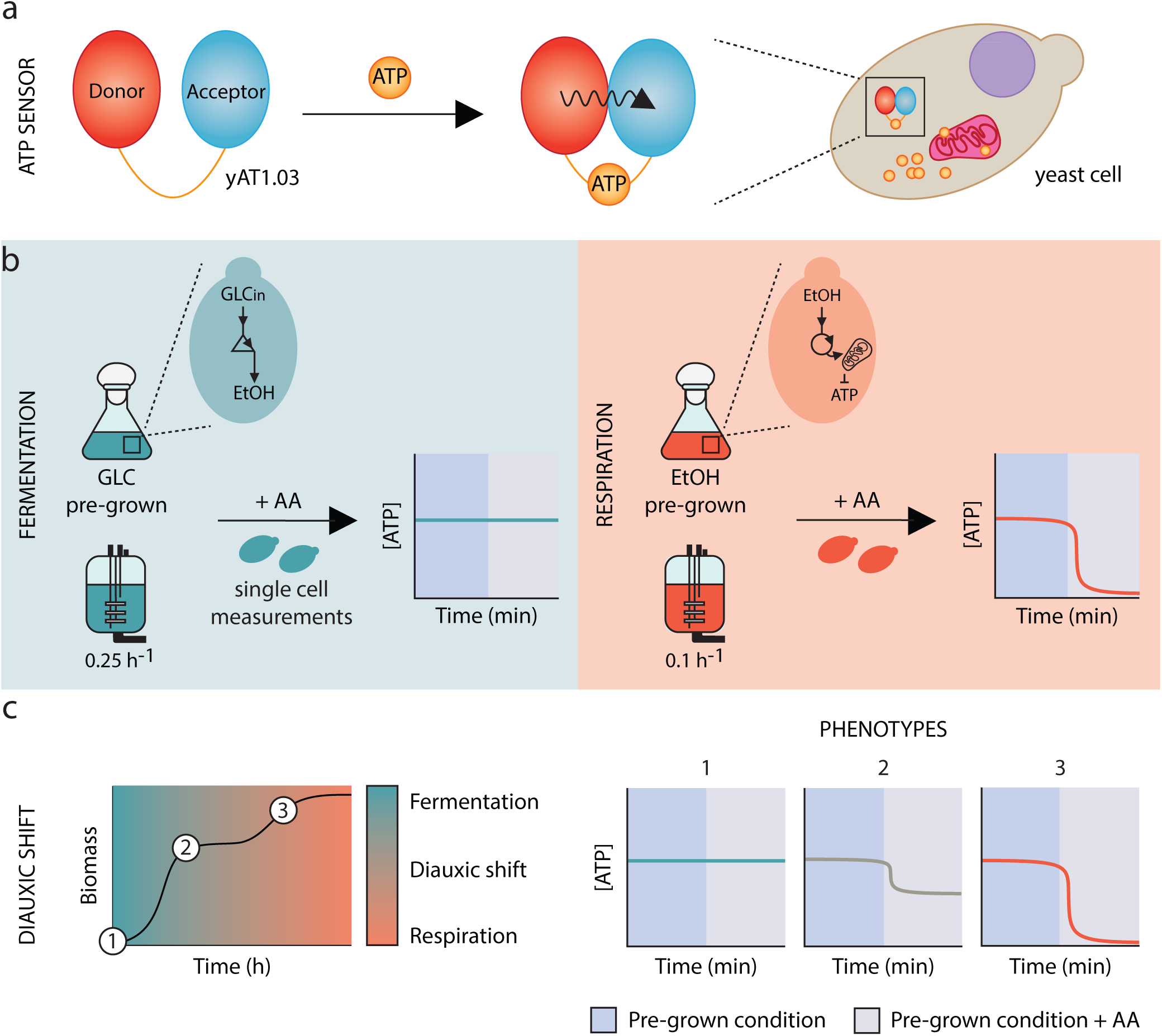

## 1 Introduction

Adenosine triphosphate (ATP), universally regarded as the main energy currency of Life, is involved in essential cell processes such as growth, intracellular transport and regulation of central carbon metabolism^1–5^. Cells rely on catabolic processes to synthesize ATP, either through respiration and proton motive force driven ATPase, or through fermentation and substrate-level phosphorylation. *Saccharomyces cerevisiae* (*S. cerevisiae*) and other Crabtree-positive yeast ferment glucose to ethanol in the presence of oxygen^6–8^. When glucose is exhausted, cells switch to ethanol respiration, generating additional ATP. This transition is known as the diauxic shift (DS) and involves extensive remodeling of the cell metabolome and proteome^9,10^. Among those is the expansion and extensive reshaping of mitochondria that creates an intricate tubular network enabling cells to grow fully respiratory^11,12^.

Proteomic changes in yeast cell makeup towards respiratory metabolism were shown to start gradually in the mitochondria, when glucose is still available^10^. It is still unclear if this gradual shift is caused by a gradual shift in each individual cell, or a population average artefact of heterogeneous but sudden shifts at the single cell level. A few studies have investigated single-cell responses of metabolism. Van Heerden used intracellular pH to follow metabolism during start up of glycolysis^13^; similarly Bagamery used mitochondrial abundance and morphology to assess metabolic states after glucose starvation and recovery^14^; Takaine measured ATP in mutants of ATP metabolism and linked it to protein aggregation^15^. None directly assessed the functional contribution of respiration versus fermentation to ATP homeostasis and the cell heterogeneity during the diauxic shift. Techniques that come closest to asses mitochondrial function are based on measuring changes in mitochondrial membrane potential and include the use of an oxygraph together with uncoupling agents in intact cells or isolated mitochondria, or the use of fluorescent dyes^16,17^. While the former method lacks single-cell resolution, the latter provides only indirect measurements (Table S1).

Here, we established a novel single-cell method that directly and quickly can distinguish between respiration or fermentation as the dominant mode of ATP production in single cells. The method relies on the expression of a Förster Resonance Energy Transfer (FRET) bio-based sensor (yAT1.03)^18^ to measure cytosolic ATP changes upon the addition of a respiration blocker (Antimycin A, AA). After being pulsed with AA, respiratory cells demonstrated a drop in ATP concentration as a result of the impairment of the electron transport chain^19^, whereas cells that ferment maintained the original ATP levels. We then applied the method to a yeast population during the diauxic shift to test for the presence of potential metabolic subpopulations.

## 2 Results

### 2.1 Induction of ATP drain in fermentative and respiratory cells after a pulse with a glucose analog

In this work we used the bio-based FRET sensor yAT1.03 to measure changes in cytosolic ATP in CEN.PK. In parallel, the strain CEN.PK + mTq2Δ11 was generated (Figure S1 a) to correct for bleedthrough of the acceptor (tdTomato) into the donnor chanel (mTq2Δ11) (Figure S1 b, c), accounting for possible differences in fluorescence emission when compared with CEN.PK + mTq2. When expressed in CEN.PK, yAT1.03 doesn’t compromise growth, however a slightly decrease in growth rate under ethanol conditions and final biomass was observed (Figure S2).

We started by testing the suitability of the ATP sensor yAT1.03 by pulsing cells with an ATP drain that should result in a decrease in cytosolic ATP in both fermentative and respiratory regimes. To do so, we submitted a population of yeast cells pre-grown on either ethanol or glucose to 2-Deoxy-D-glucose (2-DG). 2-DG is imported into the cytosol by hexose transporters and phosphorylated by hexokinase into 2-deoxy-d-glucose-6-phosphate (2-DG-6P), a step that requires one ATP molecule. The lack of the hydroxyl group in the backbone of 2-DG-6P makes it unrecognizable by glucose-6-phosphate isomerase, resulting in an blockage of glycolysis^20^. Cells grown on both carbon sources experienced a drop in ATP concentration upon 2-DG addition and no recovery was observed 8 minutes after the pulse (Figure 1 a). Quantitatively we observed a more heterogeneous response of cells grown on ethanol (Figure 1 b).

**Figure 1.**
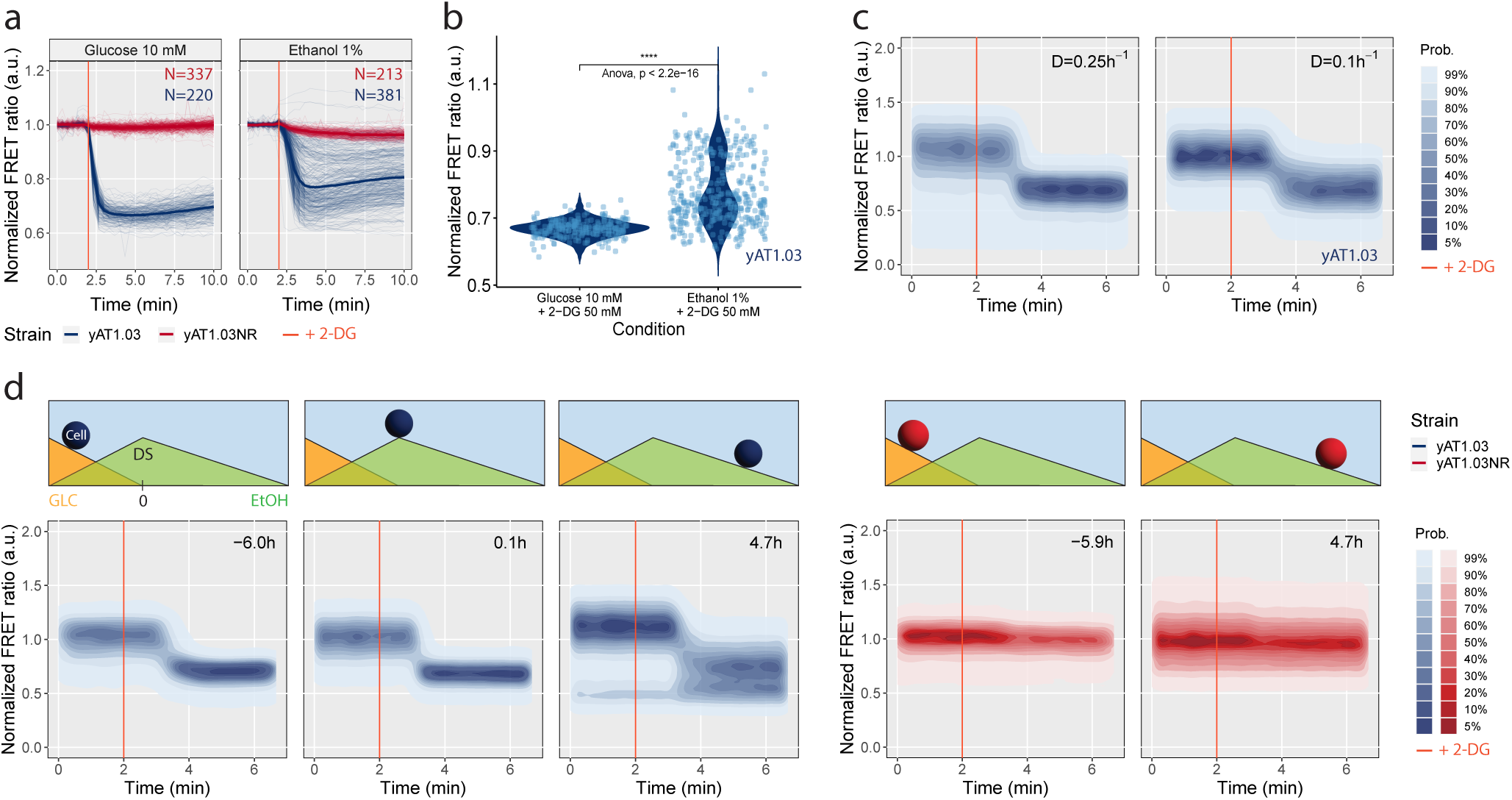
Decrease in intracellular ATP triggered by an ATP drainer measured in both metabolic state. Cells expressing the responsive (yAT1.03) and the non-responsive (yAT1.03NR) sensors were incubated in minimal media supplemented with 10 mM glucose or 1% ethanol followed by a pulse with 50 mM of 2-Deoxy-D-glucose (2-DG; concentration selected based on a dose-response curve can be found in Figure S3). The ATP dynamics were measured by fluorescent microscopy (a, b). Displayed in a), thin lines illustrate single-cell trajectories (N number of cells) and thick lines the mean FRET ratio normalized to the baseline. (b) Violin plots of the means of yAT1.03 displayed in a) measured between 3.5 and 4.5 min. The statistical tests Mann-Whitney U (p-value <0.0001) and Anova were applied to the data. (c) ATP changes measured by flow cytometry upon 2-DG addition to cells pre-grown in an aerobic controlled stepwise chemostat (D=0.1 and 0.25 *h^-^*^1^). (d) ATP dynamics measured by flow cytometry after 2-DG addition to cells expressing yAT1.03 (pre-, during and post-diauxic shift) and yAT1.03NR (pre- and post-diauxic shift). The sampling time is indicated in the upright corner of each subplot. Top diagrams illustrate the metabolic state of the cell, from growth on glucose (GLC), through the diauxic shift (DS) to growth on ethanol (EtOH).

Next, we applied the same perturbation to a culture grown in a chemostat at 0.1 and 0.25 *h^−^*^1^ dilution rates, aiming to achieve respiratory and respirofermentative metabolisms when providing the same carbon source^21^. The chemostat data shows that the cultures transitioned indeed from a respiratory to a fermentative state (ethanol production rates of 0 *mmol · g^−^*^1^ *· h^−^*^1^ at 0.09 *h^−^*^1^ and 7 *mmol · g^−^*^1^ *· h^−^*^1^ at 0.26 *h^−^*^1^, respectively) (Table 1). 2-DG addition resulted again in a drop in cytosolic ATP levels for both dilution rates (Figure 1 c, full data in Figure S4). In parallel, we monitored the glucose concentration of the samples used in the pulse experiment. Despite very low glucose concentrations, as expected, ATP levels remained responsive to 2-DG addition (Figure S5, Table S2).

**Table 1.** Chemostat cultivations show dilution rate dependent respiratory and fermentative metabolisms. Glucose grown cells at the dilution rate of 0.1 *h^−^*^1^ display a respiratory behaviour while cells grown at dilution rate of 0.25 *h^−^*^1^ display a fermentative behaviour. Data originated from biological duplicates. *Y_x/s_*is yield of biomass on glucose; *q* is specific flux with negative values meaning consumption; *c* is concentration measured in the final broth.

| Variable | D = $0.1\ h^{-1}$ | D = $0.25\ h^{-1}$ |
| --- | --- | --- |
| *Growth rate* ($h^{-1}$) | $0.09 \pm 0$ | $0.26 \pm 0$ |
| $Y_{x/s}$ ($g_{DW} \cdot g_{glucose}^{-1}$) | $0.42 \pm 0$ | $0.2 \pm 0.03$ |
| $q_{ethanol}$ ($mmol \cdot g_{DW}^{-1} \cdot h^{-1}$) | $0 \pm 0$ | $6.95 \pm 1.34$ |
| $q_{glucose}$ ($mmol \cdot g_{DW}^{-1} \cdot h^{-1}$) | $-1.25 \pm 0.01$ | $-7.06 \pm 0.94$ |
| $q_{glycerol}$ ($mmol \cdot g_{DW}^{-1} \cdot h^{-1}$) | $0 \pm 0$ | $0 \pm 0$ |
| $q_{pyruvate}$ ($mmol \cdot g_{DW}^{-1} \cdot h^{-1}$) | $0 \pm 0$ | $0 \pm 0$ |
| $q_{succinate}$ ($mmol \cdot g_{DW}^{-1} \cdot h^{-1}$) | $0 \pm 0$ | $0 \pm 0$ |
| $c_{glucose,out}$ ($mmol \cdot l^{-1}$) | $0.01 \pm 0$ | $0.09 \pm 0.01$ |
| $c_{ethanol,out}$ ($mmol \cdot l^{-1}$) | $0 \pm 0$ | $10.17 \pm 0.63$ |

Finally, we followed the response to 2-DG addition during the diauxic shift. Pre- and post-diauxic shift cells equally experienced a drop in ATP concentration after the perturbation, but again growth on ethanol appeared to give a more heterogeneous response (Figure 1 d). No response was detected for the non-responsive sensor, here used as a control.

### 2.2 Disruption of mitochondrial ATP synthesis allows to distinguish between fermentative and respiratory metabolisms

Earlier computational work had shown that yeast cells grown on glucose excess rely mostly on substrate-level phosphorylation for ATP synthesis: only 20% of total ATP production flux comes from respiration^21^. We therefore wondered if we could distinguish fully respiratory growth from respiro-fermentative growth by inhibiting respiration and monitoring the response to cytosolic ATP levels. We submitted a yeast cell population expressing yAT1.03 to Antimycin A, an inhibitor of complex III of the respiratory chain. Using both microscopy and flow cytometry, we observed a decrease in ATP levels in ethanol pre-grown cells upon AA addition, but no change in glucose pre-grown cells (microscopy data displayed in Figure 2 a, b and flow cytometry data in Figure S6).

**Figure 2.**
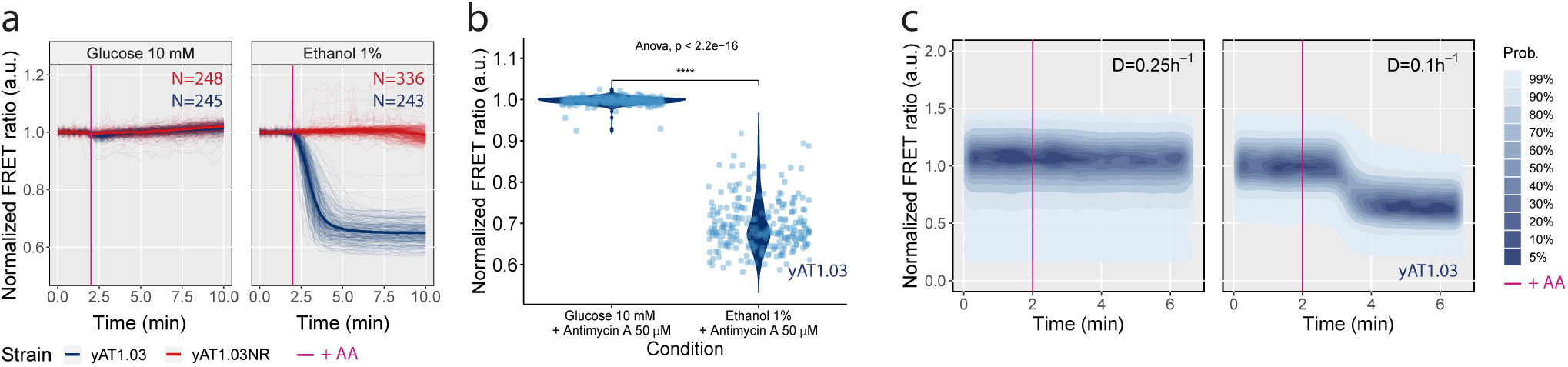
ATP dynamics upon Antimycin A addition in respiratory and fermentative cells. (a, b) Cells incubated in minimal media supplemented with 10 mM glucose or 1% ethanol were pulsed with 50 *µ*M of AA and fluorescence was followed by microscopy and flow cytometry (Figure S6) in two independent experiments. Thin lines in a) illustrate the single-cell trajectories (N number of cells) and thick lines the mean FRET ratio normalized to the baseline. (b) Violin plots of the means of yAT1.03 displayed in a) measured between 3.5 and 4.5 min. The statistical tests Mann-Whitney U (p-value <0.0001) and Anova were applied to the data. (c) ATP changes upon AA addition to cells pre-grown in an aerobic controlled stepwise chemostat (D=0.1 and 0.25 *h^-1^*).

Next, we turned to aerobic glucose-limited chemostat conditions, where *S. cerevisiae* remains respiratory at low dilution rates and becomes fermentative at high dilution rates. Cells growing at 0.1 *h^−^*^1^ suffered a drop in ATP levels after pulsed with AA, whereas no effect was observed for cells growing at 0.25 *h^−^*^1^ (Figure 2 c; full data in Figure S7). No subpopulations were observed in any of the experiments.

### 2.3 ATP dynamics during the diauxic shift reveal distinct respiratory capacities

Finally we followed the changes in fermentation versus respiration in individual cells during the diauxic shift. We sampled a glucose batch culture expressing yAT1.03 over time and pulsed cells with Antimycin A. Additional samples were collected to measure growth and metabolites concentration (Figure 3 a, b). The population-based specific rates of glucose uptake and ethanol excretion/uptake were then calculated (Figure 3 c). Altogether these data allowed us to define the time window corresponding to the diauxic shift and thus the pre- and post-DS phases. The pre-diauxic phase spanned for 20 hours, from inoculation until glucose depletion. This phase was followed by the post-diauxic shift phase characterized by ethanol consumption and a decrease in growth rate from 0.37 ± 0.05 to 0.08 ± 0 *h^−^*^1^. After being pulsed with AA, pre-diauxic shift cells showed no change in ATP FRET ratio, whereas post-diauxic shift cells experienced a decrease in the ratio (Figure 3 d). During the 8 hours that followed the diauxic shift, we observed an increasing drop in ATP levels until it reached 0.6 (mean value in Figure 3 e and min value in Figure 3 f; replicates data in Figure S8). At this point, cells are fully respiratory and have consumed 55% of the original ethanol available. No physiological subpopulations were spotted throughout the experiment, with exception of a fraction of cells with a low FRET ratio that becomes visible 2.8 h after the DS and that dominates after ethanol exhaustion (Figure S9), suggesting the presence of dead cell material. No response was observed for the non-responsive sensor after AA addition in the different stages of growth.

**Figure 3.**
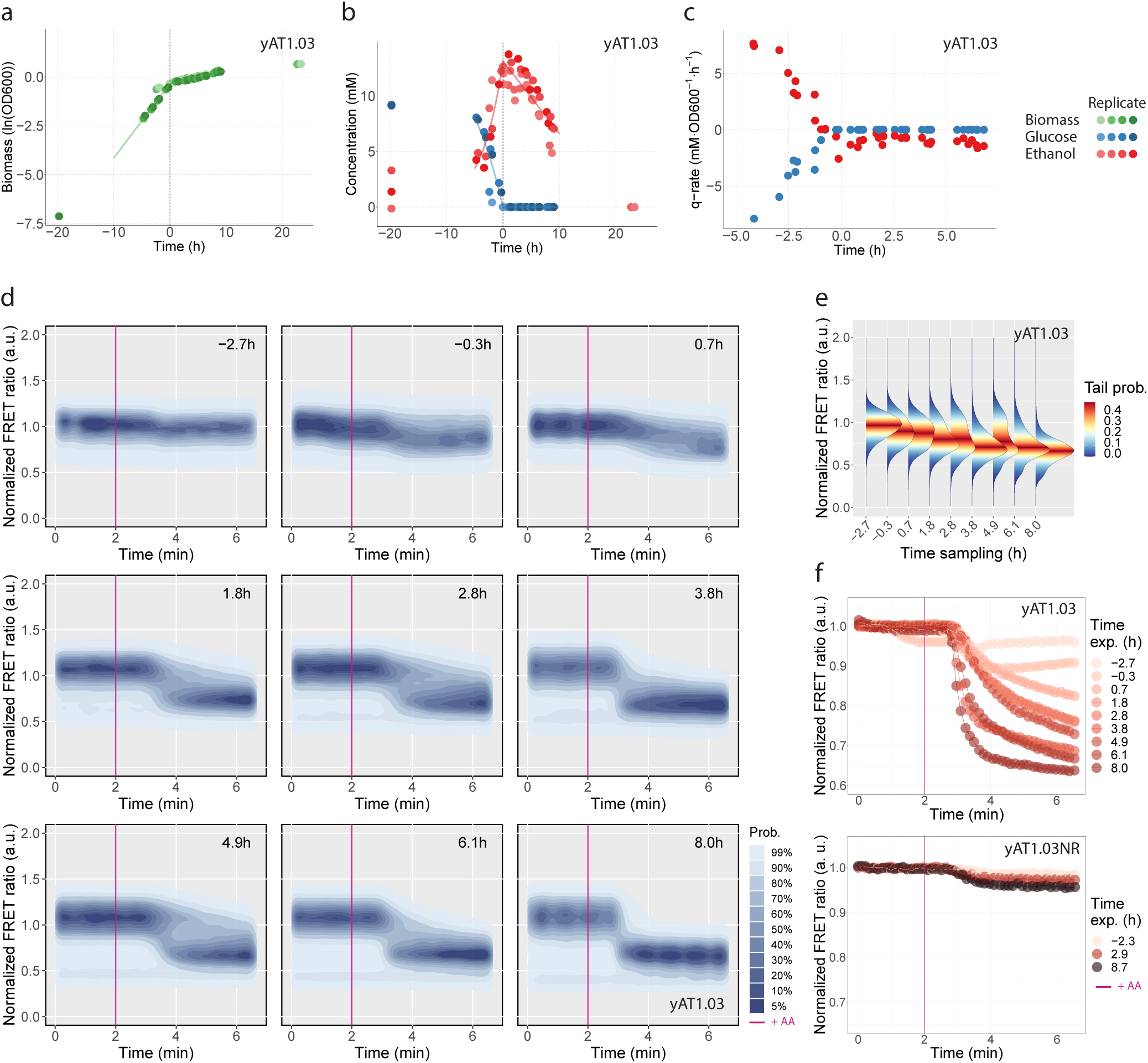
ATP dynamics during the diauxic shift reveal distinct respiratory capacities. Cells were grown in minimal media supplemented with 10 mM glucose. (a) Growth profile. (b) Glucose and ethanol concentrations. (c) Glucose and ethanol biomass specific rates. (d) Normalized ATP FRET ratio response to an AA pulse measured by flow cytometry. Data normalized to the baseline, before the pulse. The sampling time is indicated in the upright corner of each subplot. (e) Distribution plots of d) after the effect of AA becomes visible (3.33 min from the start). (f) Binned plots of the ATP FRET ratio (mean data of 50 consecutive time intervals) for the responsive (yAT1.03) and non-responsive (yAT1.03NR) sensors. Plots a), b) and c) include data of four biological replicates and plots d), e) and f) data of a representative experiment.

## 3 Discussion

In this work we developed a direct method to distinct fermentative from respiratory metabolism in *in vivo* single cells of *S. cerevisiae*. The novel method is based on the expression of the fluorescent ATP sensor yAT1.03 and short term perturbations with the respiration inhibitor Antimycin A. We validated our method in fully fermentative and respiratory metabolic regimes from batch and chemostat cultures. Finally, we used the yAT1.03 + AA method to follow the diauxic transition from glucose fermentation to ethanol respiration in a yeast cell population.

The glucose analogue 2-DG was used in a control experiment to induce expected changes in cytosolic ATP levels regardless of the carbon source provided (Figure 1). In glucose grown cells, 2-DG acts as a glucose competitor, directly disrupting ATP synthesis. In ethanol-grown cells ATP exhaustion is a parallel glycolytic pathway in cells geared towards gluconeogenesis and ATP generation in mitochondria. Considering that glycolysis is downregulated during respiration^10,22,23^, the observed ATP drop in this regime confirms the preparedness of yeast to catabolize its preferred carbon source, glucose. Metabolic anticipation has been previously demonstrated in budding yeast^24,25^. Curiously, we found a great variability in the cell response of ethanol grown cells to 2-DG. This heterogeneity could be the result of a variation in the expression of hexose transporters and hexokinase or in the nucleocytoplasmic distribution of Hxk2^26^. Cell-to-cell heterogeneity in response to unpredictably changing environments, also known as bet-hedging, is well known in yeast^14,27–29^, and is expected to be higher at lower growth rates.

Antimycin A acts by impairing the mitochondrial electron transport chain complex III (coenzyme Q: cytochrome c – oxidoreductase)^19^, compromising ATP synthesis through oxidative phosphorylation. The effect of a sudden block of respiration on the ATP concentration depends on the relative contribution of respiration to the overall ATP synthesis flux, and the flexibility of either substrate-level phosphorylation to take over ATP demand, or of ATP demand to lower in view of the limited supply. We tested both ethanol-grown cells (no option at all to scale up substrate-level phosphorylation) and cells grown fully respiratory on glucose (at a low dilution rate, little substrate to respond to a drop in ATP). In both cases, with > 90% of ATP coming from respiration and no backup from glycolysis, Antimycin A caused a drop in ATP. As expected, the ATP levels of fermenting cells remained constant after AA addition, both for glucose batch and chemostat at 0.25 *h^−^*^1^ cultures (Figure 2 a, b) despite low glucose levels in the latter case. Yeast cells growing on excess glucose rely for 80% on glycolysis to generate ATP. Moreover, glycolysis is responsive to ATP demand^13^ and under glucose excess has the ability to accelerate (which may be considered the reverse of the Pasteur effect^30^). For the chemostat sample at a D=0.25 *h^−^*^1^, respiration cannot have contributed much to ATP homeostasis as a fast response of glycolysis is ruled out by the low glucose levels under these conditions. Yet, the sharp differences observed in the ATP pattern for respiratory and fermentative cells provide a clear separation of the two metabolic phenotypes required for the method design.

After establishing the method, we applied it to a yeast cell population transitioning from glucose fermentation to ethanol respiration during diauxic growth. We found that during the lag phase following glucose exhaustion, yeast cells progressively increase their respiratory capacity (gradual increase in ATP drop). Previous works linked the increase in respiratory capacity with the expansion of the number of mitochondria and its functionality^14^. During the time course of the experiment no subpopulations were identified, indicating that all cells coordinate their response towards the diauxic shift, and the change in the slope and possibly end state of ATP (Figure 3 f and Figure S10) reflect changes in average capacity. One may expect that gradual and predictable changes in the environment should result in uniform responses, but there are notable exceptions, at least for prokaryotes^31^.

## 4 Conclusions

All together, our results show that yAT1.03 + AA is a robust method to distinguish fermentative from respiratory metabolisms in budding yeast. The method allows *in vivo* and single-cell measurements, providing a versatile platform that can be used in multiple growth setups (batch, chemostat, microfluidics) and instruments (plate reader, flow cytometer, microscope). Moreover, the short duration of the assay makes it a powerful method for quick measurements. yAT1.03 + AA brings new possibilities to study yeast metabolism under more realistic conditions often found in the biotechnology industry, such as in poorly stirring bioreactors and fermented products.

## 5 Methods

### 5.1 Strains

The prototrophic strain CEN.PK113-7D (MATa, URA3, HIS3, LEU2, TRP1, MAL2-8c, SUC2) and CEN.PK113-5D (MATa, ura3-52, HIS3, LEU2, TRP1, MAL2-8c, SUC2) were the yeast strains used in this work. The strains were used to harbor different plasmids, according the purposed of the experiment (Table 2). *E. coli* DH5-alpha was the bacterial strain selected for the cloning procedures.

**Table 2.** Yeast strains and plasmids.

| Strain background | Plasmid | Fluorescent protein(s) | Strain short name | Source plasmid | Source |
| --- | --- | --- | --- | --- | --- |
| CEN.PK113-7D | - | - | 7D | - | - |
| CEN.PK113-5D | pDRF1-GW | - | 5D | W. Frommer | Addgene #36026 |
| CEN.PK113-5D | yAT1.03 pDRF1-GW | ymTq2Δ11, tdTomato | yAT1.03 | D. Botman | Addgene #132781 |
| CEN.PK113-5D | yAT1.03R122KR126K pDRF1-GW | ymTq2Δ11, tdTomato | yAT1.03NR | D. Botman | Addgene #132782 |
| CEN.PK113-5D | tdTomato pDRF1-GW | tdTomato | tdTomato | D. Botman | in-house |
| CEN.PK113-5D | ymTq2 pDRF1-GW | ymTq2 | ymTq2 | D. Botman | Addgene #118453 |
| CEN.PK113-5D | ymTq2Δ11 pDRF1-GW | ymTq2Δ11 | ymTq2Δ11 | This work | in-house |

### 5.2 ymTq2**Δ**11 construction

The following construction was performed with the purpose of creating the exact same donor FP (lacking the last 11 amino acids) incorporated in the yAT1.03 sensor and in the correspondent non-responsive variant. pDRF1-GW ymTq2Δ11 was generated by removing the yAT.03 from pDRF1-GW yAT1.03 and ligating the opened vector with the ymTq2 lacking the last 33 nucleotides. To amplify ymTq2Δ11, the following primers were used: FW-TAAGCAGCTAGCACTAGTAAGCTTTTAA and RV-TAAGCAGCG-GCCGCTTAAGCAGCAGTAACGAATTCC, containing restriction sites for *Nhe*I and *Not*I, respectively. The PCR was performed with Phusion Hot Start II DNA Polymerase (2 U/µL) (Thermo Fisher Scientific, Scientific, Waltham, MA, USA) according the manufacture specifications. A temperature of 55*^◦^*C was used for the annealing of the primers. The PCR product was first purified using the GeneJET PCR Purification Kit (Thermo Fisher Scientific) to concentrate the DNA. A second purification step was performed in a 1% agarose gell using the GeneJET Gel Extraction Kit (Thermo Fisher Scientific) to remove the original vector used as template in the PCR reaction. pDRF1-GW yAT1.03 and ymTq2Δ11 were digested with *Nhe*I HF and *Not*I HF (enzymes from New England Biolabs, Ipswich, Massachusetts, USA) and the opened vector pDRF1-GW and insert ymTq2Δ11 purified from a 1% agarose gel using the GeneJET Gel Extraction Kit. The vector and the insert were ligated using T4 DNA Ligase (Thermo Fisher Scientific) and transformed in *E. coli* cells. The transformation was performed in chemocompetent cells according the Transformation Protocol of New England Biolabs (https://international.neb.com/protocols/2012/05/21/transformation-protocol). Positive candidates were selected for plasmid DNA isolation using the GeneJET Plasmid Miniprep Kit (Thermo Fisher Scientific). The construction pDRF1-GW ymTq2Δ11 was confirmed by DNA sequencing (https://www.macrogen-europe.com/). The new plasmid was introduced in CEN.PK113-7D by yeast transformation (performed according the protocol described by Tutucci E. *at al*.^32^). Yeast cells were selected in a plate of YNB medium -uracil Drop Out^33^ supplemented with 100 mM glucose. Fluorescence was confirmed by microscopy.

### 5.3 Plate reader growth experiments

A single colony of each strain was inoculated in 5 ml of YNB media supplemented with 100 mM glucose. Cells were diluted twice in fresh media and kept in mid-log phase until they were collected for the experiment. Cells were then washed twice in YNB supplemented with 10 mM glucose (4000 rpm at room temperature), and resuspended in the same media to an OD_600_ of 0.05. A volume of 500 *µ*L of the cell suspension was transferred to a 48 wells plate and the OD_600_ was measured in a FLUOstar Omega microplate reader every 5 minutes. Cells were kept at 30*^◦^*C and 700 rpm orbital shaking.

### 5.4 Batch growth conditions and sampling

Cells from a single colony where inoculated in YNB medium (6.8 g/L Yeast Nitrogen Base and 10.2 g/L phthalate-Phosphate buffer, pH adjusted to 5 with KOH) supplemented with 100 mM glucose and grown at 30*^◦^*C and 200 rpm to mid-log phase. Cells were washed twice in YNB medium supplemented with 10 mM glucose and grown overnight. Growth was monitored by measuring the OD_600_ of culture, using a Ultraspec 2100 pro (Amersham Biosciences, Cambridge, England). Samples for metabolites concentrations where filtered with a 0.2 *µ*m polyethersulfone (PES) filter and kept at −20*^◦^*C until further use.

### 5.5 Chemostat growth conditions and sampling

Chemostat cultures of CENPK-113-5D expressing pDRF1-GW yAT1.03 were grown at 30*^◦^*C in a 1.2 L bioreactor (Applikon, Delft, the Netherlands) with a maintained working volume of 0.5 L and a stirrer speed of 500 rpm. Media was prepared as described by Verduyn *et al.*^34^ and supplemented with 10 mM glucose as carbon source and additional 0.2 g/L of antifoam C (Sigma-Aldrich). Dissolved oxygen concentration was maintained above 40% throughout the cultivation. Steady-state was assumed after eight generation times. After the initial batch phase, indicated by rise in oxygen levels in the off-gass, pumps were turned on to maintain a dilution rate of 0.1 h*^−^*^1^ for the first sampling. The dilution rate was later increased to 0.25 h*^−^*^1^ for the second sampling. Chemostat samples were diluted with spent media, filtered with a 0.2 *µ*m PES filter, to an OD_600_ of 1 at room temperature. Samples were taken from the diluted cell suspension for flow cytometry. During the time span of the flow cytometry analysis, sub-samples from the original cell suspension were collected through a 0.2 *µ*m PES filter and stored at −20*^◦^*C for subsequent glucose concentration measurements. Cell dry weight was determined by filtering 5 mL of the culture on weighed and dried membrane filters of 0.4 *µ*m. Filters were washed with demi water and dried at 60*^◦^*C for 24 hours prior to weighing.

### 5.6 Metabolite concentration determination

Glucose concentrations from the chemostat samples were determined enzymatically with a solution of hexokinase/glucose-6-phosphate dehydrogenase (Roche) in Pipes buffer at pH 7. Measurements were performed in a FLUOstar Omega microplate reader. After addition of the enzymes, samples were incubated for 15 minutes and absorbance was measured at 340 nm. Glucose concentrations were determined by comparison of the end point measurements with a glucose calibration curve. The calibration curve was measured as described above with known glucose concentrations and in parallel with the samples. All measurements were performed in triplicate and the background absorbance was measured from samples without enzyme addition.

Samples from the batch and chemostat cultures were analysed for quantification of glucose and ethanol concentrations by High-Performance Liquid Chromatography (HPLC). Samples were thawed at 4*^◦^*C O/N and measured next day on a prominence HPLC (SHIMADZU, Kyoto, Japan) equiped with an Rezex ROA organic acid H^+^ column (Phenomex, CA, USA). A post-run analysis (LabSolutions v5.71) using calibration curves with known retention times and concentrations was performed to retrieve the metabolites concentrations. Four (responsive and non-responsive sensors) or two (single FPs) independent cultures were included in the analysis.

### 5.7 Flow cytometry

The ATP levels were followed in a flow cytometer CytoFLEX S (Beckman Coulter, Brea, USA) using the software CytExpert (v2.4.0.28). The flow rate was set to 10 *µ*L/min and constrained to 2000 events/sec. A violet layser was used in combination with the RFP-405nm (610/20) and CFP-405nm (470/20) filters. A tube containing 920 *µ*L of the cell suspension was placed in the machine and the baseline measured for 2 minutes. 100 *µ*L of 0.5 mM AA diluted in 5% ethanol was added to the cells and mixed with a pipette. The response was measured for additional 5 minutes. Samples were processed using the slow speed running mode. In between samples we performed a first 1 min wash with absolute ethanol (to remove possible AA residues in the machine) and a second 2 min wash with demi water (to wash the ethanol) using the fast speed running mode. In order to correct the fluorescence of tdtomato for ymTq2Δ11 bleedtrough, we measured the fluorescence in the RFP-405nm and CFP-405nm channels of the strain ymTq2Δ11 at different time points of the diauxic shift and applied the average. A minimum number of 10.000 events was considered.

### 5.8 Microscopy

The yeast strain CENPK.113-5D expressing yAT1.03 or yAT1.03R122KR126K were used in the following experiments. Cells were grown in YNB media supplemented with 1% ethanol or 100 mM glucose at 200 rpm and 30*^◦^*C to mid-log phase. Next, cells were transferred to a Attofluor cell chamber (Thermofisher Scientific, Waltham, MA, USA), containing a ConA coated coverslip (prepared according Hansen *et al.*^35^), and incubated at 30*^◦^*C for 15 minutes, following two washing steps in 900 *µ*L of fresh media. Ethanol pre-grown cells were washed in the same media and glucose pre-grown cells on YNB 10 mM glucose. Cells were placed in a Nikon Ti-eclipse widefield fluorescence microscope (Nikon, Minato, Tokio, Japan) at 30°C for imaging. A 2 minutes baseline was recorded before the addition of 100 *µ*L of Antimycin A (final concentration 50 *µ*M in ethanol 0.5%) or 2-DG (final concentration 10-100 mM). After the pulse, the response was followed for additional 8 minutes. Cells were imaged every 20 seconds and FRET was recorded using a TuCam system (Andor, Belfast, Northern Ireland) equipped with 2 Andor Zyla 5.5 sCMOS cameras (Andor). A 438/24 nm excitation filter, a 483/32 nm donor emission filter and a 593/40 nm acceptor emission filter (552 nm long-pass dichroic filter) (Semrock, Lake Forest, IL, USA) were used. The light intensity was adjusted to 7.4 in a SOLA 6-LCR-SB power source (Lumencor, Beaverton, OR, USA). Cell segmentation was performed using an in-house macro in FiJi (NIH, Bethesda, MD, USA). First, image drift across channels was corrected using the 2D image stabilizer plugin TurboReg^36^ developed by P. Thevenaz *et al.* and background correction was performed. Next, cells were segmented using the Weka Segmentation plugin^37^ developed by Arganda-Carreras *et al.* and the brigthest channel (CFP) as reference. The mean fluorescence of each cell was calculated for each fluorescent channel. A filter based on size and round shape was applied to remove cell aggregates (min size of 2000 or 3000 pixels depending on the dataset and max size of 1200 pixels; 0.6-1 roundness).

### 5.9 Data analysis

R Core Team (2013) was used for data analysis and visualization. R: A language and environment for statistical computing. R Foundation for Statistical Computing, Vienna, Austria. ISBN 3-900051-07-0, URL http://www.R-project.org/. The R package flowCore^38^ was used for cell gating and filtering. The fluorescence of tdtomato was corrected for ymTq2Δ11 bleedtrough (0.774 of ymTq2Δ11 fluorescence) and the normalized FRET ratio was calculated as RFP/CFP (or tdTomato/ymT2Δ11) regularized to the baseline (before the addition of the compound, and considered until 1.3 and 2 minutes for the flow cytometry and microscopy analyses, respectively). The distribution plots of the ATP dynamics after AA addition include the data of the second half of the experiment (3.33 min from the start). The FRET ratio illustrated in the binned plot was calculated using the discretize function of the R package arules^39^ and 50 breaks. To calculate the ATP consumption rate we applied a breakpoint analysis using segmented regressions^40^. To determine the biomass specific rates, a rolling window with 6 constitutive measurements was used. The biomass specific rate was calculated by multiplying the growth rate by the metabolite yield on biomass. The biomass and metabolites concentrations experimental data were fitted by splitting it into two subsets, before and after glucose exhaustion, and fitting each of the new subsets independently. The substrate yield on biomass was calculated by linear regression on the biomass concentration (OD600) versus metabolite concentration (mM). The growth rate was calculated by applying linear regressions on the natural logarithm of time (h) versus biomass concentration (OD600).

## Acknowledgements

The CEN.PK113-7D strain was kindly provided by P. Kötter, Euroscarf, Frankfurt. We acknowledge the financial support from the Dutch Research Council (NWO) (Project numbers 731.016.001 and ENPPS.LIFT.019.005). 731.016.001 is a public private-partnership with DSM and ENPPS.LIFT.019.005 with Chr Hansen. We thank Dennis Botman for technical support.

## Author contribution

L.L. wrote the manuscript, conceptualize the idea and together with J.B. designed the experiments. L.L., J.B., D.J. and V.E.Z. performed the experiments. L.L., J.B. and C.M. analysed the data. B.T. supervised the work.

## Abbreviations

AA: Antimycin A
ATP: Adenosine Triphosphate
DS: Diauxic Shift
*E. coli*: *Escherichia coli*
EtOH: Ethanol
FRET: Förster Resonance Energy Transfer
GLC: Glucose
HPLC: High-Performance Liquid Chromatography
*S. cerevisiae*: *Saccharomyces cerevisiae*
OD: Optical Density
PES: Polyethersulfone
2-DG: 2-Deoxy-D-glucose

## References

1 . Jorgelindo da Veiga Moreira, Sabine Peres, Jean-Marc Steyaert, Erwan Bigan, Loïc Paulevé, Marcel Levy Nogueira, and Laurent Schwartz. Cell cycle progression is regulated by intertwined redox oscillators. Theoretical Biology and Medical Modelling, 12(1), May 2015.

2 . Wei-Hsiang Lin and Christine Jacobs-Wagner. Connecting single-cell ATP dynamics to overflow metabolism, cell growth, and the cell cycle in escherichia coli. Current Biology, 32(18):3911–3924.e4, September 2022.

3 . Kaspar P Locher. Mechanistic diversity in ATP-binding cassette (ABC) transporters. Nature Structural &amp Molecular Biology, 23(6):487–493, June 2016.

4 . Christer Larsson, Inga lill Phlman, and Lena Gustafsson. The importance of ATP as a regulator of glycolytic flux inSaccharomyces cerevisiae. Yeast, 16(9):797–809, 2000.

5 . Sindhuja Sridharan, Nils Kurzawa, Thilo Werner, Ina Günthner, Dominic Helm, Wolfgang Huber, Marcus Bantscheff, and Mikhail M. Savitski. Proteome-wide solubility and thermal stability profiling reveals distinct regulatory roles for ATP. Nature Communications, 10(1), March 2019.

6 . Johannes P van Dijken, Ruud A Weusthuis, and Jack T Pronk. Kinetics of growth and sugar consumption in yeasts. Antonie van leeuwenhoek, 63:343–352, 1993.

7 . Carl Malina, Rosemary Yu, Johan Björkeroth, Eduard J. Kerkhoven, and Jens Nielsen. Adaptations in metabolism and protein translation give rise to the crabtree effect in yeast. Proceedings of the National Academy of Sciences, 118(51), December 2021.

8 . Julius Battjes, Chrats Melkonian, Sebastián N. Mendoza, Auke Haver, Kosai Al-Nakeeb, Anna Koza, Lars Schrubbers, Marijke Wagner, Ahmad A. Zeidan, Douwe Molenaar, and Bas Teusink. Ethanol-lactate transition of lachancea thermotolerans is linked to nitrogen metabolism. Food Microbiology, 110:104167, April 2023.

9 . Guillermo G Zampar, Anne Kümmel, Jennifer Ewald, Stefan Jol, Bastian Niebel, Paola Picotti, Ruedi Aebersold, Uwe Sauer, Nicola Zamboni, and Matthias Heinemann. Temporal system-level organization of the switch from glycolytic to gluconeogenic operation in yeast. Molecular Systems Biology, 9(1):651, January 2013.

10 . Francesca Di Bartolomeo, Carl Malina, Kate Campbell, Maurizio Mormino, Johannes Fuchs, Egor Vorontsov, Claes M. Gustafsson, and Jens Nielsen. Absolute yeast mitochondrial proteome quantification reveals trade-off between biosynthesis and energy generation during diauxic shift. Proceedings of the National Academy of Sciences, 117(13):7524–7535, March 2020.

11 . Alexander Egner, Stefan Jakobs, and Stefan W. Hell. Fast 100-nm resolution three-dimensional microscope reveals structural plasticity of mitochondria in live yeast. Proceedings of the National Academy of Sciences, 99(6):3370–3375, March 2002.

12 . Carl Malina, Christer Larsson, and Jens Nielsen. Yeast mitochondria: an overview of mitochondrial biology and the potential of mitochondrial systems biology. FEMS Yeast Research, 18(5), April 2018.

13 . Johan H. van Heerden, Frank J. Bruggeman, and Bas Teusink. Multi-tasking of biosynthetic and energetic functions of glycolysis explained by supply and demand logic. BioEssays, 37(1):34–45, October 2014.

14 . Laura E. Bagamery, Quincey A. Justman, Ethan C. Garner, and Andrew W. Murray. A putative bet-hedging strategy buffers budding yeast against environmental instability. Current Biology, 30(23):4563–4578.e4, December 2020.

15 . Masak Takaine, Hiromi Imamura, and Satoshi Yoshida. High and stable ATP levels prevent aberrant intracellular protein aggregation in yeast. eLife, 11, April 2022.

16 . Andrea Volejníková, Jana Hlousková, Karel Sigler, and Alena Pichová. Vital mitochondrial functions show profound changes during yeast culture ageing. FEMS Yeast Research, 13(1):7–15, February 2013.

17 . Magdalena Kwolek-Mirek and Renata Zadrag-Tecza. Comparison of methods used for assessing the viability and vitality of yeast cells. FEMS Yeast Research, pages n/a–n/a, September 2014.

18 . Dennis Botman, Johan H. van Heerden, and Bas Teusink. An improved ATP FRET sensor for yeast shows heterogeneity during nutrient transitions. ACS Sensors, 5(3):814–822, February 2020.

19 . Xiuquan Ma, Mingzhi Jin, Yu Cai, Hongguang Xia, Kai Long, Junli Liu, Qiang Yu, and Junying Yuan. Mitochondrial electron transport chain complex III is required for antimycin a to inhibit autophagy. Chemistry &amp Biology, 18(11):1474–1481, November 2011.

20 . Martin C. Schmidt and Allyson F. O’Donnell. ‘sugarcoating’ 2-deoxyglucose: mechanisms that suppress its toxic effects. Current Genetics, 67(1):107–114, November 2020.

21 . Ibrahim E. Elsemman, Angelica Rodriguez Prado, Pranas Grigaitis, Manuel Garcia Albornoz, Victoria Harman, Stephen W. Holman, Johan van Heerden, Frank J. Bruggeman, Mark M. M. Bisschops, Nikolaus Sonnenschein, Simon Hubbard, Rob Beynon, Pascale Daran-Lapujade, Jens Nielsen, and Bas Teusink. Whole-cell modeling in yeast predicts compartment-specific proteome constraints that drive metabolic strategies. Nature Communications, 13(1), February 2022.

22 . Aaron PALOMINO, Pilar HERRERO, and Fernando MORENO. Rgt1, a glucose sensing transcription factor, is required for transcriptional repression of the iHXK2/i gene in isaccharomyces cerevisiae/i. Biochemical Journal, 388(2):697–703, May 2005.

23 . Roeland Costenoble, Paola Picotti, Lukas Reiter, Robert Stallmach, Matthias Heinemann, Uwe Sauer, and Ruedi Aebersold. Comprehensive quantitative analysis of central carbon and amino-acid metabolism in isaccharomyces cerevisiae/i under multiple conditions by targeted proteomics. Molecular Systems Biology, 7(1):464, January 2011.

24 . Amir Mitchell, Gal H. Romano, Bella Groisman, Avihu Yona, Erez Dekel, Martin Kupiec, Orna Dahan, and Yitzhak Pilpel. Adaptive prediction of environmental changes by microorganisms. Nature, 460(7252):220–224, June 2009.

25 . Riddhiman Dhar, Rudolf Sägesser, Christian Weikert, and Andreas Wagner. Yeast adapts to a changing stressful environment by evolving cross-protection and anticipatory gene regulation. Molecular Biology and Evolution, 30(3):573–588, November 2012.

26 . Deifilia Ahuatzi, Alberto Riera, Rafael Peláez, Pilar Herrero, and Fernando Moreno. Hxk2 regulates the phosphorylation state of mig1 and therefore its nucleocytoplasmic distribution. Journal of Biological Chemistry, 282(7):4485–4493, February 2007.

27 . Sasha F. Levy, Naomi Ziv, and Mark L. Siegal. Bet hedging in yeast by heterogeneous, age-correlated expression of a stress protectant. PLoS Biology, 10(5):e1001325, May 2012.

28 . Aaron M. New, Bram Cerulus, Sander K. Govers, Gemma Perez-Samper, Bo Zhu, Sarah Boogmans, Joao B. Xavier, and Kevin J. Verstrepen. Different levels of catabolite repression optimize growth in stable and variable environments. PLoS Biology, 12(1):e1001764, January 2014.

29 . Jue Wang, Esha Atolia, Bo Hua, Yonatan Savir, Renan Escalante-Chong, and Michael Springer. Natural variation in preparation for nutrient depletion reveals a cost–benefit tradeoff. PLOS Biology, 13(1):e1002041, January 2015.

30 . J. A. Den Hollander, K. Ugurbil, T. R. Brown, M. Bednar, C. Redfield, and R. G. Shulman. Studies of anaerobic and aerobic glycolysis in saccharomyces cerevisiae. Biochemistry, 25(1):203–211, January 1986.

31 . Ana Solopova, Jordi van Gestel, Franz J. Weissing, Herwig Bachmann, Bas Teusink, Jan Kok, and Oscar P. Kuipers. Bet-hedging during bacterial diauxic shift. Proceedings of the National Academy of Sciences, 111(20):7427–7432, May 2014.

32 . Evelina Tutucci, Maria Vera, and Robert H. Singer. Single-mRNA detection in living s. cerevisiae using a re-engineered MS2 system. Nature Protocols, 13(10):2268–2296, September 2018.

33. Dropout mix. Cold Spring Harbor Protocols, 2006(1):pdb.rec8585, June 2006.

34 . C. Verduyn, E. Postma, W. A. Scheffers, and J. P. van Dijken. Physiology of saccharomyces cerevisiae in anaerobic glucose-limited chemostat culturesx. Journal of General Microbiology, 136(3):395–403, March 1990.

35 . Anders S Hansen, Nan Hao, and Erin K O’Shea. High-throughput microfluidics to control and measure signaling dynamics in single yeast cells. Nature Protocols, 10(8):1181–1197, July 2015.

36 . P. Thevenaz, U.E. Ruttimann, and M. Unser. A pyramid approach to subpixel registration based on intensity. IEEE Transactions on Image Processing, 7(1):27–41, 1998.

37 . Ignacio Arganda-Carreras, Verena Kaynig, Curtis Rueden, Kevin W Eliceiri, Johannes Schindelin, Albert Cardona, and H Sebastian Seung. Trainable weka segmentation: a machine learning tool for microscopy pixel classification. Bioinformatics, 33(15):2424–2426, March 2017.

38 . B Ellis, Perry Haaland, Florian Hahne, Nolwenn Le Meur, Nishant Gopalakrishnan, Josef Spidlen, Mike Jiang, and Greg Finak. flowCore: flowCore: Basic structures for flow cytometry data, 2022. R package version 2.8.0.

39 . Michael Hahsler, Christian Buchta, Bettina Gruen, and Kurt Hornik. arules: Mining Association Rules and Frequent Itemsets, 2023. R package version 1.7-6.

40 . Vito M.R. Muggeo. Interval estimation for the breakpoint in segmented regression: a smoothed score-based approach. Australian &amp New Zealand Journal of Statistics, 59(3):311–322, September 2017.

