## Supplementary material for "A fast method to distinguish between fermentative and respiratory metabolisms in single yeast cells": S1

#### List of Figures

S1 **Fluorescence emitted by CEN.PK113-5D + mTq2Δ11 in the red and blue channels and respective ratio during a diauxic shift experiment.** (a) Yeast cells expressing ymTq2 or ymTq2Δ11 were pre-grown on glucose and visualized using widefield microscopy. (b) Fluorescence of CEN.PK113-5D + mTq2Δ11 was measured in the red and blue channels by flow cytometry and the respective ratio calculated (c). Despite the increase in fluorescence observed for both channels, the contribuition was the same in both channels, resulting in a constant ratio over time (0 h corresponding to the diauxic shift). Plot b) combines replicate A and B, while plot c) shows the replicates separately. . . . 1

S2 **Growth profiles of the yeast strains used in this work.** Cells were growing in YNB media supplemented with 10 mM glucose. The prototrophic strain CEN.PK113-7D and the strain CEN.PK113-5D + pDRF1 were used as controls. Data include two biological replicates and two technical replicates per experiment. . . . 2

S3 **2-DG dose response in CEN.PK113-5D + yAT1.03.** Cells pre-grown on 100 mM glucose were washed and incubated in 10 mM glucose followed by addition of 2-DG (min 2). Fluorescence was measured by widefield microscopy, allowing to track individual responses, here represented by a thin line. The FRET ratio was normalized to the baseline (dark blue area) and plotted over time. . . . 3

S4 **Response of cells grown in a chemostat to a 2-DG pulse.** Cells expressing yAT1.03 were grown in a chemostat at 0.1 and 0.25 $h^{-1}$ followed by addition of 50 mM 2-DG (min 2). FRET ratio measured by flow cytometry and normalized to the baseline. Data is originated from two biological replicates (A and B) and replicate B was selected to be displayed in the main manuscript. . . . 4

S5 **Glucose depletion in chemostat samples.** The glucose concentration of the chemostat samples taken for the pulse experiment was monitored through time to confirm that a residual amount would still be present at the moment of the experiment. . . . 5

S6 **ATP dynamics upon Antimycin A addition to respiratory and fermentative cells.** Cells were incubated in YNB media supplemented with 10 mM glucose or 1% ethanol and pulsed with 50 $\mu$M of AA. Fluorescence was measured by flow cytometry. . . . 6

S7 **Response of cells grown in a chemostat to an Antimycin A pulse.** Cells expressing yAT1.03 were grown in a chemostat at 0.1 and 0.25 $h^{-1}$ followed by addition of AA (min 2). FRET ratio measured by flow cytometry and normalized to the baseline. Data is originated from two biological replicates (A and B) and replicate B was selected to be displayed in the main manuscript. . . . 7

S8 **ATP dynamics upon Antimycin A addition to yeast cells pre- during and post-diauxic shift.** Binned plots of the normalized ATP FRET ratio (mean data of 50 consecutive data points) of yAT1.03 and yAT1.03 after AA adition. Plots include data of four biological replicates (A, B, C and D). Replicate A is included in the main manuscript. . . . 8

S9 **ATP dynamics post-diauxic shift shows the existence of a low-ATP subpopulation.** Starved cells display a low normalized FRET ratio when compared with cells grown on glucose or ethanol and are unable to respond to an Antimycin A pulse. The time on the top right corner represents the sampling time relative to the diauxic shift. . . . . . . . . . . . 9
S10 **Relationship between maximum ATP synthesis inhibition and delta FRET ratio.** (a) Zoom in on the normalized FRET ratio throughout the diauxic shift experiment (Figure 3 f) between 2.7 and 3.5 minutes and respective slopes generated by linear regression. (b) Slopes generated by a breakpoint analysis using segmented regressions for the dataset shown in a) and time experiment 8 h. (c) Delta change of the normalized FRET ratio vs maximum absolute slope. Slopes were calculated according the method displayed in b). . . . 10

#### List of Tables

S1 **Current methods to assess yeast respiration.** (*) Refers to intact cells or isolated mitochondria. (**) Limited to single-cell RNA-sequencing. . . . . . . . . . . . . . . . . . . . . 11
S2 **Depletion of glucose in chemostat samples.** The glucose concentrations in the chemostat samples used in the pulse experiments was followed through time. The mean glucose concentrations per reactor, dilution rates and time points are here represented. . . . . . . . . 12

The fluorescent protein ymTq2 was modified by removing the last 11 amino acids (ymTq2Δ11) in order to accurately mimic the acceptor protein present in the FRET sensor yAT1.03 used in this work and previously described by Botman *et al.*[1]. The fluorescence of the new ymTq2Δ11 cyan variant was tested by microscopy (Figure S1 a). To correct the flow cytometry data for the bleedtrought of the acceptor FP (mTq2Δ11) into the donnor FP (tdTomato), we measured the fluorescence of the strain CEN.PK113-5D + mTq2Δ11 in the blue and red channels during the diauxic shift (Figure S1 b). No differences were observed in the fluorescence ratio Red/Blue over the course of the diauxic shift experiment (Figure S1 c) and the mean value was used for correction proposes of the tdTomato fluorescence.

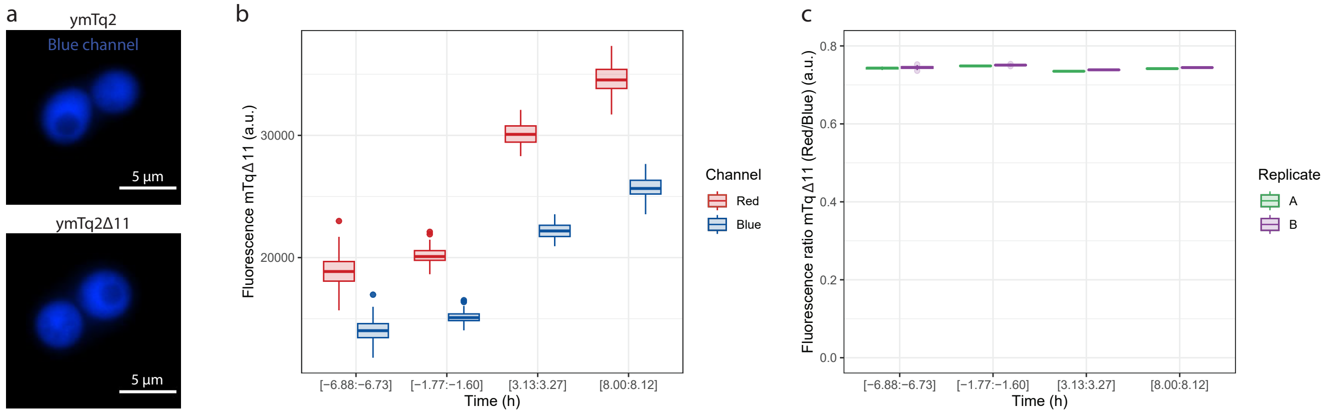


**Figure S1. Fluorescence emitted by CEN.PK113-5D + mTq2Δ11 in the red and blue channels and respective ratio during a diauxic shift experiment.** (a) Yeast cells expressing ymTq2 or ymTq2Δ11 were pre-grown on glucose and visualized using widefield microscopy. (b) Fluorescence of CEN.PK113-5D + mTq2Δ11 was measured in the red and blue channels by flow cytometry and the respective ratio calculated (c). Despite the increase in fluorescence observed for both channels, the contribuition was the same in both channels, resulting in a constant ratio over time (0 h corresponding to the diauxic shift). Plot b) combines replicate A and B, while plot c) shows the replicates separately.

The growth profiles of the strains described in the present work were followed over time (Figure S2). We observed a similar growth rate on glucose across strains, however slightly differences were found for cell growing on ethanol, with the prototrophic strain CEN.PK113-7D and the strain expressing the empty vector pDRF1 displaying the highest growth rate and maximum biomass reached.

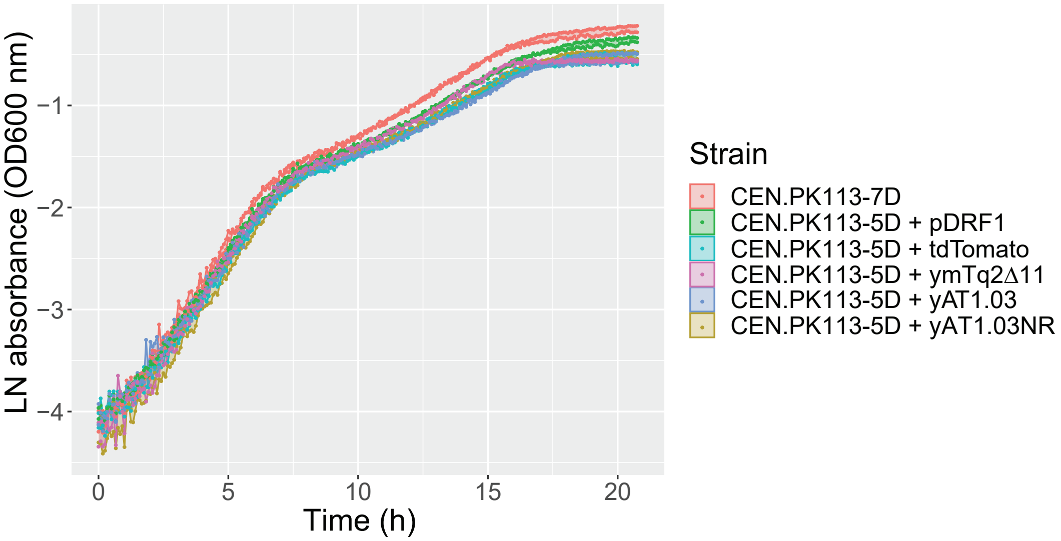


**Figure S2. Growth profiles of the yeast strains used in this work.** Cells were growing in YNB media supplemented with 10 mM glucose. The prototrophic strain CEN.PK113-7D and the strain CEN.PK113-5D + pDRF1 were used as controls. Data include two biological replicates and two technical replicates per experiment.

We started by measuring the response of CEN.PK113-5D + yAT1.03 to 10 mM 2-DG (same equimolar concentration to the glucose available), as described by Botman *et al.*[1]. However, we found that this concentration was insufficient to fully block ATP synthesis through glycolysis (Figure S3). As a result, we selected a concentration of 2-DG of 50 mM for this study.

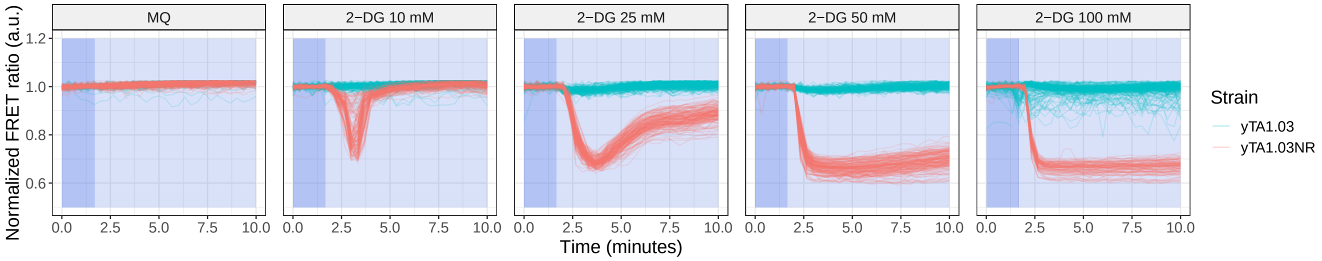


**Figure S3. 2-DG dose response in CEN.PK113-5D + yAT1.03.** Cells pre-grown on 100 mM glucose were washed and incubated in 10 mM glucose followed by addition of 2-DG (min 2). Fluorescence was measured by widefield microscopy, allowing to track individual responses, here represented by a thin line. The FRET ratio was normalized to the baseline (dark blue area) and plotted over time.

ATP dynamics of two independent chemostat cultures grown in fermentative or respiratory regimes (D=0.25 and 0.1 $h^{-1}$, respectively) after the addition of 2-DG (Figure S4).

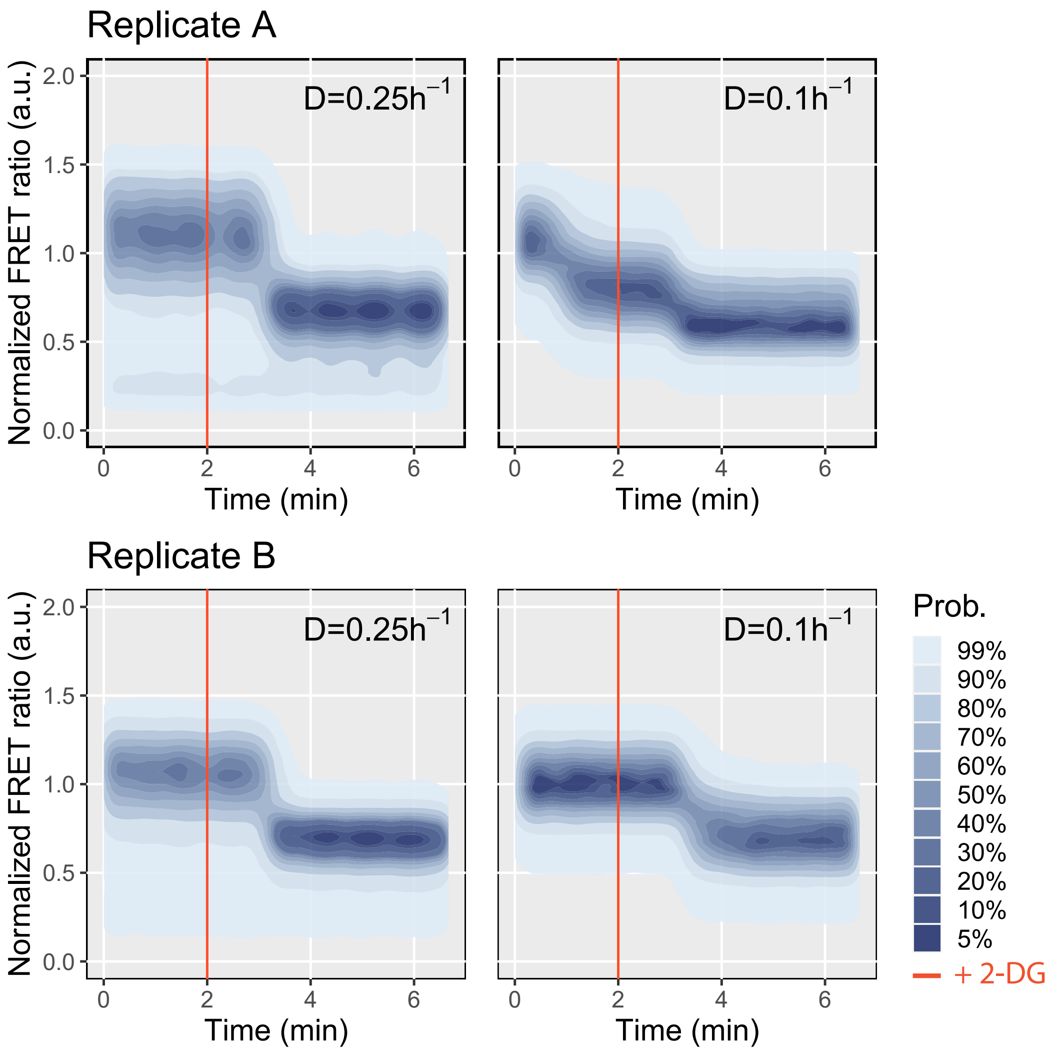


**Figure S4. Response of cells grown in a chemostat to a 2-DG pulse.** Cells expressing yAT1.03 were grown in a chemostat at 0.1 and 0.25 $h^{-1}$ followed by addition of 50 mM 2-DG (min 2). FRET ratio measured by flow cytometry and normalized to the baseline. Data is originated from two biological replicates (A and B) and replicate B was selected to be displayed in the main manuscript.

We monitored the glucose concentration over time of the sampled chemostat broth used in the pulse experiment performed in the flow cytometer (2-DG and AA) (Figure S5) (Table S2). Data show that in both cases there was glucose present during the pulse experiment. Glucose concentrations ranged between 10 and 5.5 µM and between 67 and 50 µM for the 0.1 and 0.25 $h^{-1}$ dilution rates cultures, respectively.

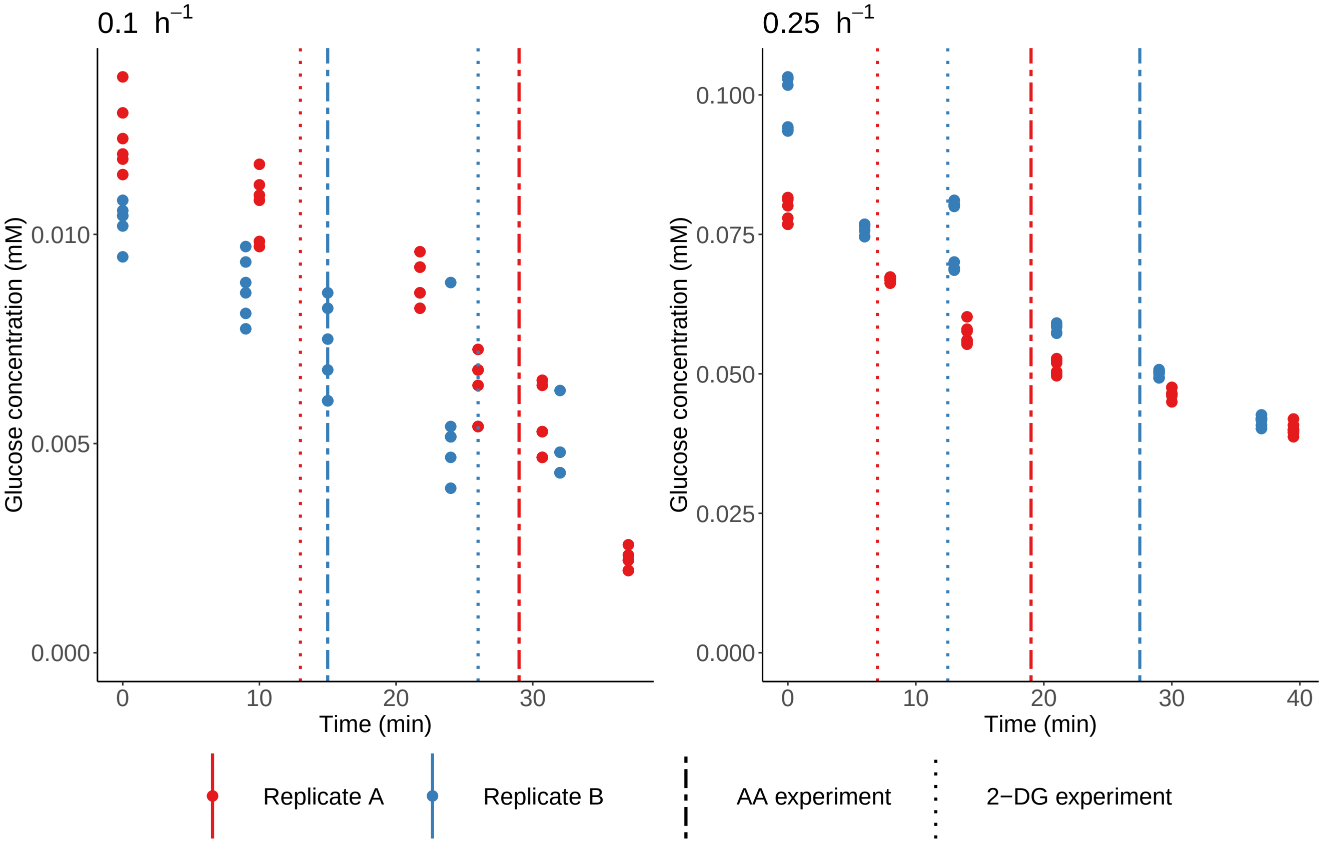


**Figure S5. Glucose depletion in chemostat samples.** The glucose concentration of the chemostat samples taken for the pulse experiment was monitored through time to confirm that a residual amount would still be present at the moment of the experiment.

We confirmed the response of yeast cells to Antimycin A addition previously obtained by microscopy by monitoring the ATP dynamics through flow cytometry under the same growth conditions (Figure S6).

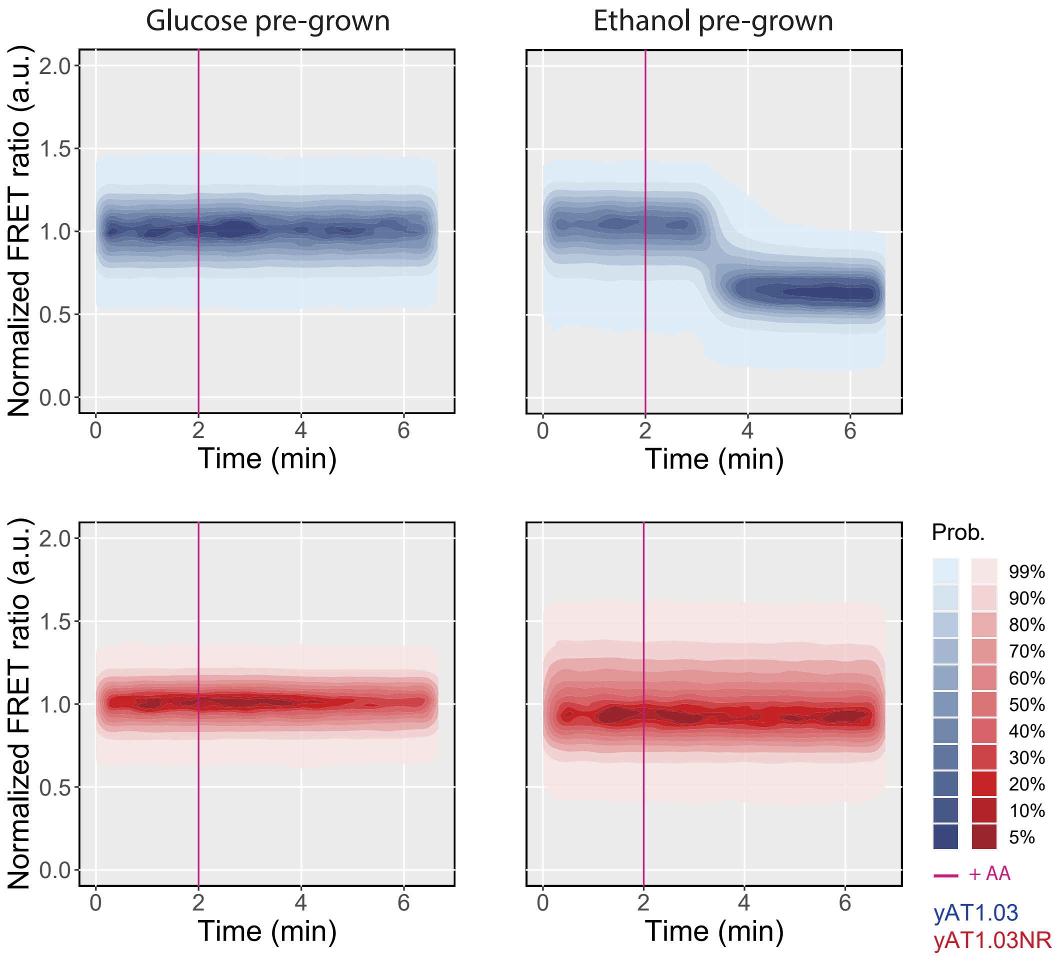


**Figure S6. ATP dynamics upon Antimycin A addition to respiratory and fermentative cells.** Cells were incubated in YNB media supplemented with 10 mM glucose or 1% ethanol and pulsed with 50 $\mu$M of AA. Fluorescence was measured by flow cytometry.

ATP dynamics of two independent chemostat cultures grown in fermentative or respiratory regimes (D=0.25 and 0.1 $h^{-1}$, respectively) after the addition of AA (Figure S7).

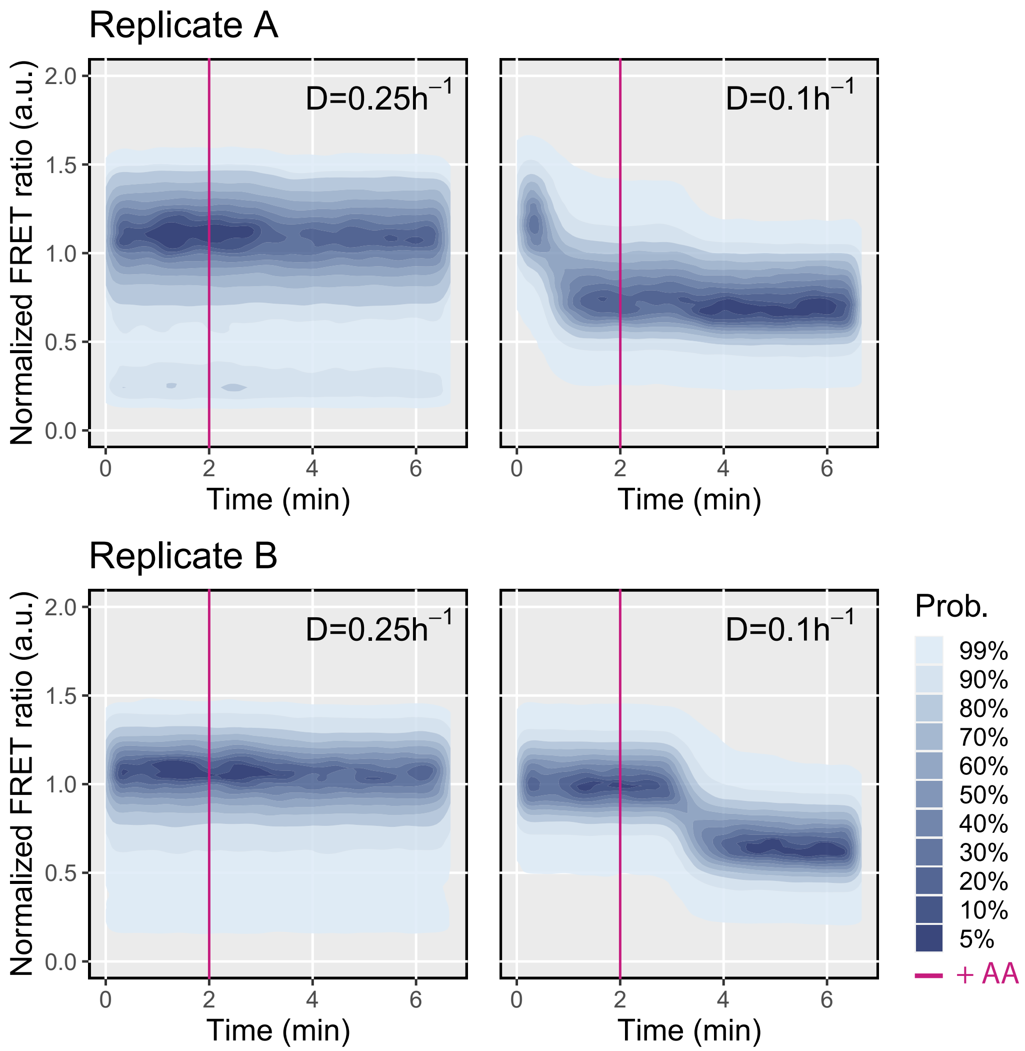


**Figure S7. Response of cells grown in a chemostat to an Antimycin A pulse.** Cells expressing yAT1.03 were grown in a chemostat at 0.1 and 0.25 $h^{-1}$ followed by addition of AA (min 2). FRET ratio measured by flow cytometry and normalized to the baseline. Data is originated from two biological replicates (A and B) and replicate B was selected to be displayed in the main manuscript.

Summary data of cells response to Antimycin A in the different stages of batch growth (glucose, diauxic shift and ethanol) (Figure S8).

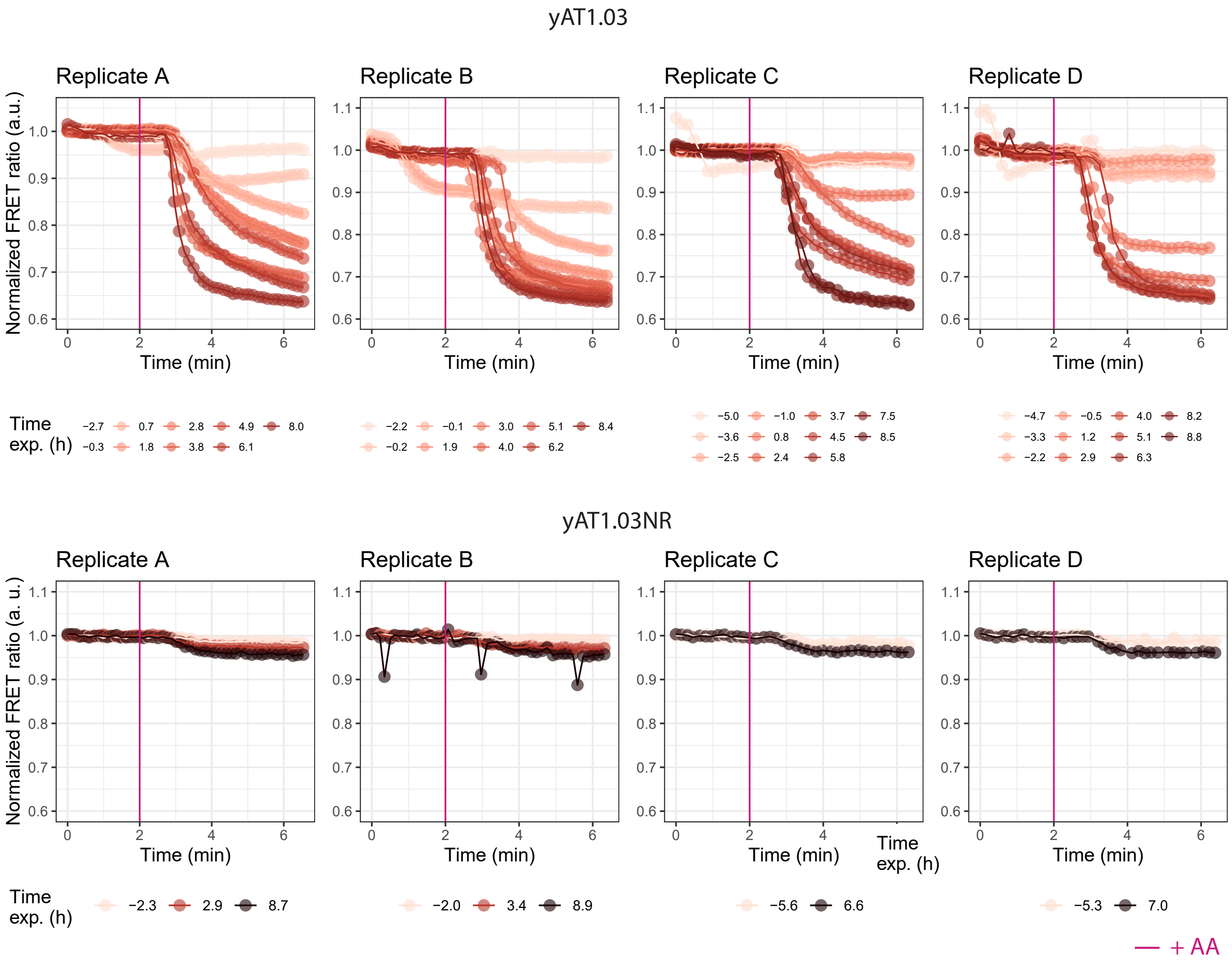


**Figure S8.** **ATP dynamics upon Antimycin A addition to yeast cells pre- during and post-diauxic shift.** Binned plots of the normalized ATP FRET ratio (mean data of 50 consecutive data points) of yAT1.03 and yAT1.03 after AA adition. Plots include data of four biological replicates (A, B, C and D). Replicate A is included in the main manuscript.

Throughout the batch experiments we consistently observed an increase in the abundance of a subpopulation with a low normalized FRET ratio (Figure S9, replicate A Figure 3 d). In the absence of carbon source, this subpopulation represents the majority of the cells.

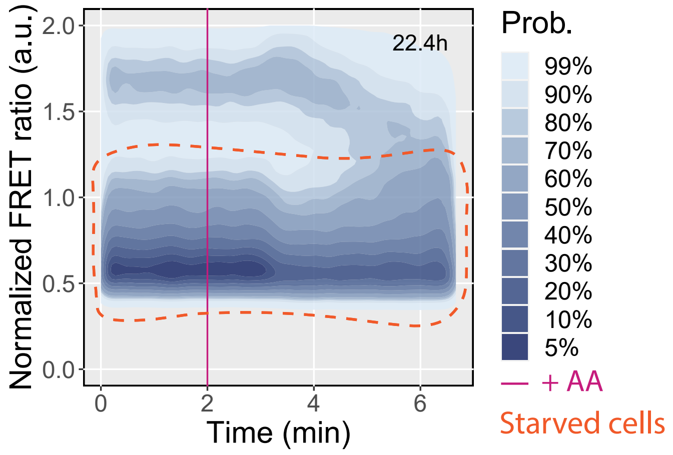


**Figure S9. ATP dynamics post-diauxic shift shows the existence of a low-ATP subpopulation.** Starved cells display a low normalized FRET ratio when compared with cells grown on glucose or ethanol and are unable to respond to an Antimycin A pulse. The time on the top right corner represents the sampling time relative to the diauxic shift.

We hypothesised that a higher speed in ATP consumption in response to Antimycin A would positively correlate with the changes in FRET ratio. To asses this, we measured the rate of ATP synthesis inhibition across the diauxic shift experiment and compared it with the difference in normalized FRET ratio before and after the addition of the compound. However we found a strong suggestion that a faster decrease in normalized FRET ratio (indicated by a higher absolute slope) leads to a lower normalized FRET ratio, this was not always the case (Figure S10).

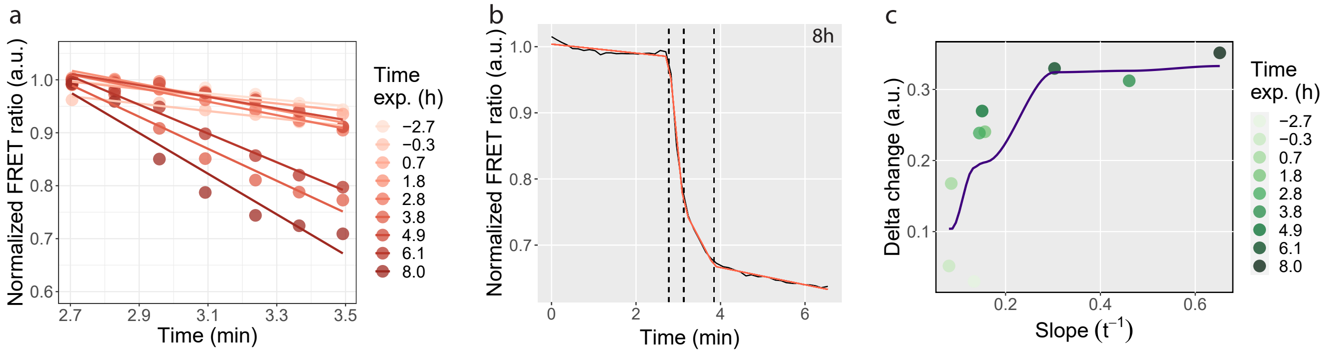


**Figure S10. Relationship between maximum ATP synthesis inhibition and delta FRET ratio.** (a) Zoom in on the normalized FRET ratio throughout the diauxic shift experiment (Figure 3 f) between 2.7 and 3.5 minutes and respective slopes generated by linear regression. (b) Slopes generated by a breakpoint analysis using segmented regressions for the dataset shown in a) and time experiment 8 h. (c) Delta change of the normalized FRET ratio vs maximum absolute slope. Slopes were calculated according the method displayed in b).

Different methods are available to measure fermentation and respiratory capacity in budding yeast (Table S1). The yAT1.03 + AA assay here developed offers for the first time a quick method to measure single cell responses to ATP dynamics, allowing to distinguish between the two metabolic regimes.

**Table S1. Current methods to assess yeast respiration.** (*) Refers to intact cells or isolated mitochondria. (**) Limited to single-cell RNA-sequencing.

| Method | Resolution | Single cell | Method | Destructive | Type | Time assay | Number of cells | Reference |
|---|---|---|---|---|---|---|---|---|
| Oxygraph | Quantitative | No | *In vivo*/ *In vitro* | Yes/No* | Direct | Few min | 4E7 | Simonovik *et al.*[2] |
| Fluorescent dyes | Semi-quantitative | Yes | *In vivo* | No | Direct | 30-45 min | 1E4 - 7E6 | Chacko *et al.*[3] |
| Omics | Quantitative | Yes**/No | *In vitro* | Yes | Indirect | Weeks | 2E7 | Di Bartolomeo *et al.*[4] |
| yAT1.03 + AA | Quantitative | Yes | *In vivo* | No | Direct | Few min | 1E4 - 7E6 | This work |

**Table S2. Depletion of glucose in chemostat samples.** The glucose concentrations in the chemostat samples used in the pulse experiments was followed through time. The mean glucose concentrations per reactor, dilution rates and time points are here represented.

| Dilution rate ($h^{-1}$) | Reactor | Time (min) | [Glucose] (μM) |
|---|---|---|---|
| 0.1 | A | 0 | 12.4 |
| 0.1 | A | 10 | 10.7 |
| 0.1 | A | 21.75 | 8.9 |
| 0.1 | A | 26 | 6.3 |
| 0.1 | A | 30.7 | 5.5 |
| 0.1 | A | 37 | 2.2 |
| 0.1 | B | 0 | 10.3 |
| 0.1 | B | 9 | 8.7 |
| 0.1 | B | 15 | 7.2 |
| 0.1 | B | 24 | 5.5 |
| 0.1 | B | 32 | 4.8 |
| 0.25 | A | 0 | 79.1 |
| 0.25 | A | 8 | 67.0 |
| 0.25 | A | 14 | 57.1 |
| 0.25 | A | 21 | 51.2 |
| 0.25 | A | 30 | 46.5 |
| 0.25 | A | 39.5 | 40.1 |
| 0.25 | B | 0 | 98.3 |
| 0.25 | B | 6 | 75.8 |
| 0.25 | B | 13 | 74.8 |
| 0.25 | B | 21 | 58.3 |
| 0.25 | B | 29 | 50.0 |
| 0.25 | B | 37 | 41.5 |

### References


1. Dennis Botman, Johan H. van Heerden, and Bas Teusink. An improved ATP FRET sensor for yeast shows heterogeneity during nutrient transitions. *ACS Sensors*, 5(3):814–822, February 2020.
2. B Simonovik and E Gnaiger. A mitochondrial reference assay for o2k high-resolution respirometry using freeze-dried baker's yeast. *Mitochondrial Physiology Network*, 15, 2017.
3. Leeba Chacko and Vaishnavi Ananthanarayanan. Quantification of mitochondrial dynamics in fission yeast. *BIO-PROTOCOL*, 9(23), 2019.
4. Francesca Di Bartolomeo, Carl Malina, Kate Campbell, Maurizio Mormino, Johannes Fuchs, Egor Vorontsov, Claes M. Gustafsson, and Jens Nielsen. Absolute yeast mitochondrial proteome quantification reveals trade-off between biosynthesis and energy generation during diauxic shift. *Proceedings of the National Academy of Sciences*, 117(13):7524–7535, March 2020.
